# Enhanced *in vivo* gene knockout with undetectable off-targets using multiplexed Cas12a sgRNAs

**DOI:** 10.1101/2024.11.26.625385

**Authors:** Fillip Port, Martha A. Buhmann, Jun Zhou, Mona Stricker, Alexander Vaughan-Brown, Ann-Christin Michalsen, Eva Roßmanith, Amélie Pöltl, Lena Großkurth, Julia Huber, Laura B. Menendez Kury, Bea Weberbauer, Maria Hübl, Florian Heigwer, Michael Boutros

## Abstract

CRISPR nuclease-mediated gene knock-out is limited by suboptimal sgRNAs, inaccessible target sites, and silent mutations. Here, we present a Cas12a-based system that targets each gene with four sgRNAs to overcome these limitations, using *Drosophila* as a tractable *in vivo* model. We show that multiplexed sgRNAs act synergistically to create deletions between target sites, substantially increasing the fraction of loss-of-function mutations. To systematically assess off-target effects, we developed a novel screening assay that visualizes CRISPR-induced chromosomal alterations in living animals. This enabled comprehensive screening of more than 2000 sgRNAs clustered in 525 quadruple arrays across 21 megabases of genomic DNA, revealing remarkably high on-target activity (100%, 82/82) and undetectable off-target cutting (0%, 0/443). Quantitative side-by-side comparisons with a current Cas9-based system targeting over 100 genes demonstrates that multiplexed Cas12a-mediated gene targeting achieves superior performance and reveals phenotypes missed by established methods. This highly efficient and specific system provides a framework for reliable functional genomics studies across diverse organisms.

## Introduction

CRISPR gene editing has significantly accelerated the functional annotation of genomes (1–3). Despite its transformative impact, several factors limit the efficiency of gene disruption through CRISPR nucleases. For example, suboptimal activity of sgRNAs or Cas proteins can impede the induction of DNA breaks and chromatin state and genetic variation can render target sites inaccessible. Moreover, repair of DNA double strand breaks (DSBs) typically results in a spectrum of mutations that includes undesired repair outcomes retaining gene function, such as small in-frame insertions or deletions (indels). As a result, CRISPR editing often produces genetic mosaics - populations of cells carrying different mutations or maintaining wild-type sequences. This severely compromises the sensitivity and robustness of functional gene discovery with CRISPR in multicellular organisms and cultured cells, presents a hurdle for the development of genetic control strategies for disease vectors, and limits its potential therapeutic application in cell and gene therapy.

A promising strategy to overcome these limitations is the simultaneous targeting of genes with multiple sgRNAs (4–6). Here, additional sgRNAs can compensate for those that fail to mediate the desired outcome. However, for Cas9, the most widely used CRISPR nuclease, creating multiplexed arrays of more than two sgRNAs remains technically challenging due to their large size and repetitive nature (4,6). Most current large-scale Cas9 sgRNA libraries thus employ only one or two sgRNAs per gene (7–12). In contrast, Cas12a nuclease is guided solely by small crRNAs (hereafter sgRNAs) and can autonomously process compact arrays into individual sgRNAs (13). These properties allow Cas12a sgRNA arrays to be encoded on commercially available oligonucleotides, substantially facilitating generation of expression plasmids at scale. This has been harnessed for multiplex gene editing and transcriptome engineering in various contexts (14–18), but whether higher-order multiplexing of sgRNAs targeting the same gene enhances the efficiency of gene disruption *in vivo* remains unclear.

A key challenge of CRISPR gene editing is the occurrence of off-target mutations. These arise when Cas nucleases cleave sites with imperfect homology to the sgRNA (19,20). This concern is particularly acute for multiplexed editing, as the chance of unintended modifications increases with each additional sgRNA. Although computational tools can predict some off-target sites, their reliability is limited by poorly understood sgRNA-specific effects and sequence differences between reference and experimental genomes. However, methods for the unbiased, experimental detection of off-target effects in the genome of most animals are limited. Methods for the direct detection of induced DNA breaks have been described, but are not available for most species due to the absence of specialized tools like antibodies binding specific DSB repair proteins (21,22). Whole genome sequencing is broadly available, but its application for comprehensive off-target analysis across many sgRNAs is limited by cost. Moreover, since CRISPR-induced mutations and natural genetic variation are often indistinguishable, inferring causal relationships about off-target effects from sequencing data remains non-trivial (23,24). Consequently, the off-target activity of CRISPR systems in many organisms remains unclear, as many studies evaluate activity at only a very small fraction of the genome or with a small number of sgRNAs or do not test for off-target cleavage at all.

Here, we describe a highly efficient gene targeting system that combines an enhanced variant of Cas12a with quadruple sgRNA arrays. When targeting sites in parallel, sgRNAs can act synergistically to induce deletions between target sites, generating mutations that are more likely to disrupt gene function. Direct comparisons with established Cas9-based methods demonstrate superior knockout efficiency of this system across diverse tissues and target genes. To comprehensively evaluate the specificity of multiplexed editing, we developed a novel assay that visualizes induced loss-of-heterozygosity (LOH) in living animals. This approach enabled an unbiased assessment of nuclease activity with over 2,000 sgRNAs in 525 arrays across 21 megabases of the genome, revealing that Cas12a editing maintains high specificity even when using multiple sgRNAs simultaneously.

## Results

### Synergistic action of quadruple Cas12a^+^ sgRNA arrays

We set out to implement and evaluate multiplexed Cas12a gene targeting in the tractable *Drosophila* model system. The short generation time of this organism enables comparisons across many conditions and replicates and the ability to encode nucleases and sgRNAs on stable, single copy transgenes in defined genomic landing sites eliminates variations in delivery efficiency or expression level as confounding factors of experimental results. We previously showed that a variant of *Lachnospiraceae bacterium* Cas12a with a D156R mutation (hereafter Cas12a^+^) functions with high efficiency in *Drosophila* and can process large sgRNA arrays (25). To determine the optimal number of sgRNAs targeting the same gene, we developed a reporter assay using genomic insertions containing one to six copies of a sgRNA target sequence (Fig. 1a). Gene editing with ubiquitously expressed Cas12a^+^ and single or multiplexed sgRNA target sites generated distinct editing outcomes in somatic cells (Fig. 1b) or the germline (Fig. 1c). Editing at single target sites primarily produced small indels, while targeting multiple sites progressively increased the frequency of larger deletions between sites (Fig. 1b, c). This highlights that sgRNA multiplexing offers benefits beyond functional redundancy, as sgRNAs act synergistically to induce larger deletions, which are particularly likely to constitute loss-of-function alleles.

**Figure 1:**
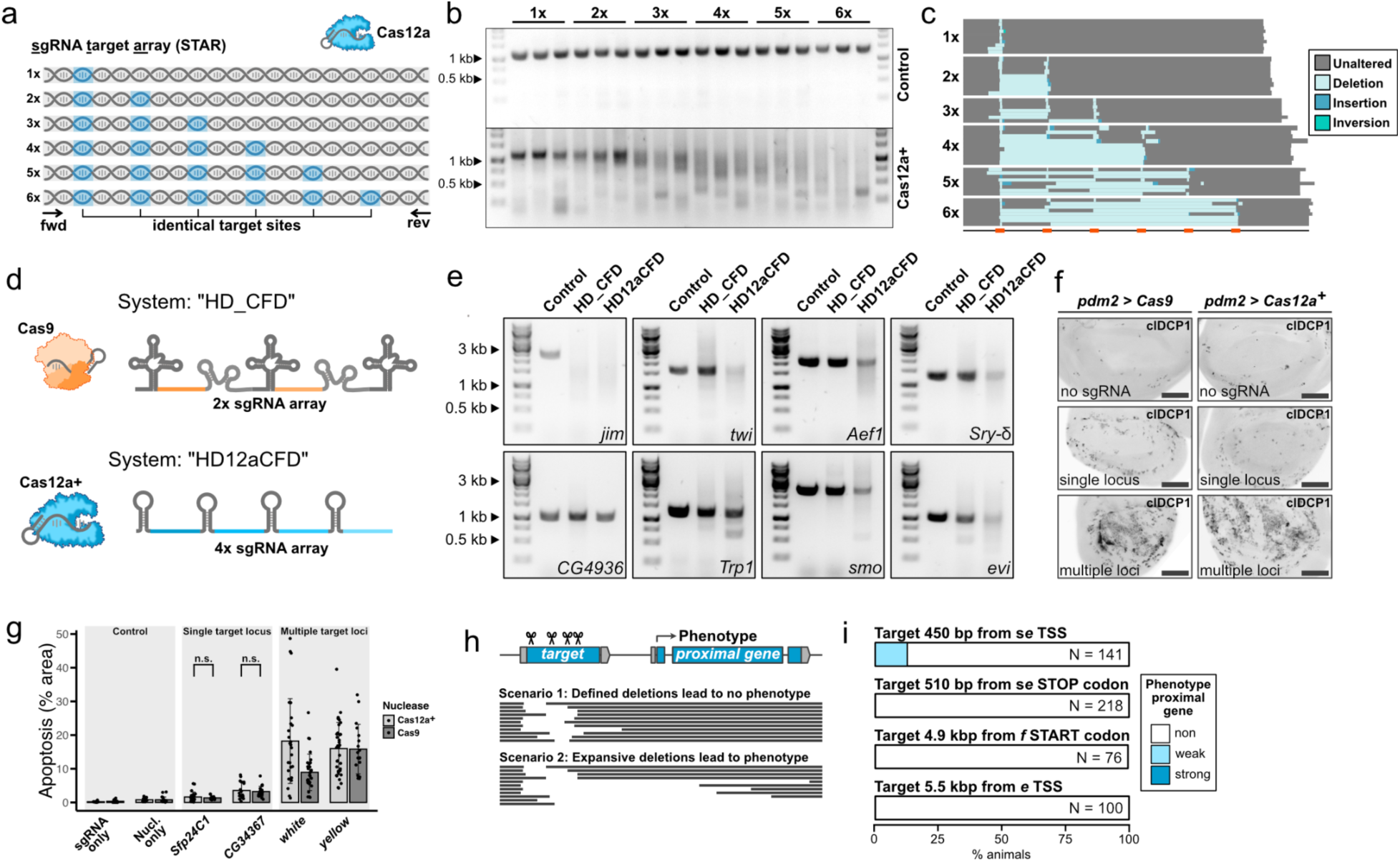
Multiplex Cas12a targeting leads to large deletions restricted to the target locus. (**a**) sgRNA target array (STAR) reporter design for assessing multiplexed CRISPR editing outcomes. It consists of a non-coding sequence containing different copy numbers of sgRNA target sites that are targeted by Cas12a^+^/sgRNA complexes *in vivo*. (**b**) Bulk editing outcomes assessed by PCR analysis of genomic DNA isolated from whole adult flies. In contrast to samples without nuclease (control; upper panel), targeting of the reporter results in amplicons of various sizes. Additional target sites produce more diverse editing outcomes, showing increased smaller amplicons and decreased amplicons of original length. Samples from three individual animals are shown for each genotype. (**c**) Individual alleles of the STAR reporter were isolated by germline transmission and identified by sequencing. Multiplexing of target sites results in frequent deletions between target sites. (**d**) Overview of the two systems used throughout most of this study. The HD_CFD system uses Cas9 nuclease and two sgRNA per gene, which are expressed from a tRNA vector, as described in (10). The HD12aCFD system uses Cas12a^+^ and an array of four sgRNA per gene. (**e**) Bulk editing outcomes after mutagenesis of endogenous genes. The target locus was amplified by PCR using identical conditions and amplicons were visualized by agarose gel electrophoresis. Control samples express sgRNAs targeting another locus. CRISPR mutagenesis frequently leads to a diversification of PCR amplicons, an effect that is notably stronger with HD12aCFD for 6 of the 8 tested target genes. (**f**) Representative images of wing imaginal discs stained for apoptotic cells with anti-cleaved *Drosophila* caspase 1 (clDCP1). Single locus sgRNAs target *CG34367*, while multi gene sgRNAs target *yellow*, which is present on all three chromosomes (see Supplementary Data Fig. 1). Scale bar: 40 µm. (**g**) Quantification of clDCP1-positive apoptotic area in the *pdm2-Gal4* expression domain. Points are individual samples, columns represent the mean and error bars the standard deviation. There is no significant (n.s.) difference whether gene targeting is performed with two sgRNAs and Cas9 or four sgRNA and Cas12a^+^ (Welch Two Sample T-Test). (**h**) Schematic of the experimental setup to detect large, expansive deletions, which are difficult to detect by PCR based assays due to deletion of a primer binding site. Using proximal genes with visible phenotypes upon knockout allows detection though phenotyping. (**i**) Results from experiments as depicted in (h). Phenotypes indicative of expansive deletions are only detected in one line with target sites in very close proximity, suggesting deletions are largely defined to the target locus. Phenotypes are presented in Extendend Data Fig.2.

We selected an intermediate number of four sgRNAs per gene for in-depth characterization to balance the benefits of multiplexing against potential drawbacks, which include excessive DNA damage and off-target effects. This choice is further supported by recent evidence that arrays consisting of more than 4 sgRNAs are no longer optimally processed by Cas12a (5). We refer to our genetically encoded quadruple sgRNA system as HD12aCFD, following on from our earlier described HD_CFD Cas9 library, which expresses two sgRNAs per gene (10) (Fig. 1d). To evaluate whether mutagenesis of endogenous genes with HD12aCFD sgRNA arrays and Cas12a^+^ also results in a high proportion of larger deletions, we assayed the integrity of the target locus after mutagenesis and compared it to animals in which the same gene was targeted with Cas9 and HD_CFD sgRNA lines (Fig. 1e). Mutagenesis with HD12aCFD sgRNA arrays resulted in a marked decrease in full length amplicons from 6 out of 8 target loci, an effect that was only observed at 1 locus following HD_CFD mediated gene targeting (Fig. 1e). These experiments reveal that the HD12aCFD system produces a diverse set of repair outcomes dominated by large deletions, a feature expected to facilitate gene inactivation.

### Multiplexed gene targeting is well tolerated and spatially restricted

We then investigated potential adverse effects of using several sgRNAs to target the same gene. Since DSBs are detrimental to cell fitness, one concern is that inducing multiple breaks simultaneously could overwhelm the DNA damage response, leading to cell death (26–28). To assess this risk, we quantified the level of apoptosis following gene targeting with multiple sgRNAs. DSBs in either of two non-essential genes with Cas9 or Cas12a^+^ caused a moderate increase in apoptotic cells in the wing disc epithelium, with comparable effects using either nuclease with two or four sgRNAs (Fig. 1f,g). In contrast, targeting genes present on multiple chromosomes resulted in a marked increase in apoptotic cells, irrespective of sgRNAs number, and produced severe adult phenotypes (Fig. 1f,g, Supplementary Data Fig. 1). These results indicate that while multiple DSBs in close proximity are well tolerated, induction of DNA breaks across different chromosomes is highly toxic.

Next, we tested whether multiplexed gene editing induces mutations in genes other than the intended target, so called off-target effects. We first focused on genes in direct proximity to the target gene, as large deletions extending beyond the target locus could lead to the disruption of neighboring genes. Such events would confound experimental results and would not be detected in PCR-based assays, as they would delete a primer binding site. Therefore, we directly tested for this possibility by analyzing four sgRNA arrays targeting genes located in direct vicinity, or intronic, to genes with well described visible phenotypes (Fig. 1f, Supplementary Data Fig. 2). Gene targeting resulted in a diverse set of large deletions at the target locus, further corroborating the results described above (Supplementary Data Fig. 2). Lines with target sites in close proximity (500-5000 bp) to the *sepia* (*se*), *ebony*, or *forked* genes yielded no phenotypes indicative of disruption of these adjacent genes (Fig. 1g and Supplementary Data Fig. 2). However, when targeting a site just 450 bp upstream of the *se* transcriptional start site, we detected a loss or reduction of *se* function in a small subset of flies (13%, N=141) with low penetrance. Collectively, these results show that multiplexed Cas12a^+^ gene targeting produces deletions that rarely extend beyond the target locus, with measurable effects on neighboring functional elements occurring only when these are in immediate proximity to the target site.

### Large-scale, unbiased off-target screening reveals high specificity

We next investigated whether multiplexed Cas12a^+^ editing induces off-target effects distant from the target locus. A meaningful evaluation of the specificity of a new gene editing system requires testing of many sgRNA throughout large segments of the genome, but suitable methods for this task have not been described in most organisms. We therefore aimed to establish an high-throughput assay for unbiased detection of CRISPR nuclease activity across megabases of genomic DNA *in vivo*. We reasoned that visualizing loss of heterozygosity (LOH), a common repair outcome of CRISPR-induced chromosome breaks in diverse systems ranging from yeast to humans (29–32), would be a suitable readout. In *Drosophila*, LOH occurs primarily through mitotic recombination (33,34) (Fig. 2a, b) and can be visualized with a heterozygous dose-sensitive reporter, such as a transgene encoding a fluorescent protein (Fig.2b). In animals with one reporter inserted on chromosome arm 2L and another on 3R, targeting Cas12a^+^ to 2L induces LOH specifically on 2L but not 3R (Fig. 2c, d). Similarly, targeting 3R with a sgRNA array induced LOH only on that chromosome arm.

**Figure 2:**
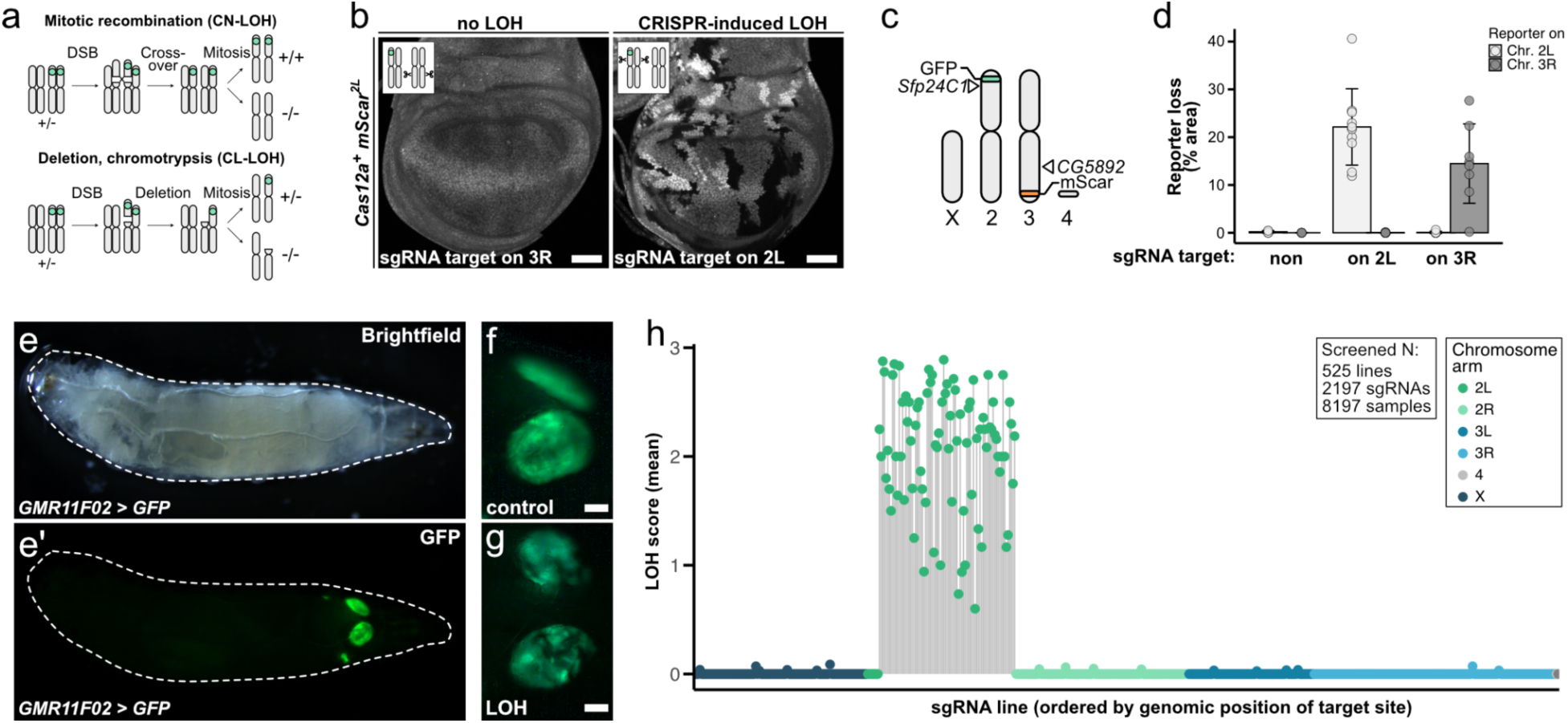
High-throughput screening reveals high on-target and undetectable off-target activity of multiplexed Cas12a^+^ editing. (**a**) Schematic illustrating mechanisms of loss of heterozygosity (LOH). Crossover between homologous chromosomes results in mitotic recombination and copy number neutral (CN) LOH, leading to homozygous daughter cells (+/+ or −/−) from heterozygous parent cells (+/−). Large chromosomal deletions or chromotrypsis lead to copy number loss (CL) LOH. (**b**) CRISPR-induced LOH in larval wing imaginal discs. Tissues express Cas12a^+^, sgRNA arrays, and have a heterozygous *mScarlet* (*mScar*) transgene inserted on chromosome arm 2L. Left panel: sgRNAs targeting a different chromosome arm (3R) do not result in LOH. Right panel: sgRNAs targeting a locus between the *mScar* transgene and the centromere result in frequent LOH resulting in loss or gain of *mScar* expression. Scale bar: 50 µm. (**c**) Schematic of the dual-reporter genotype used in (d). Two different fluorophore transgenes (*GFP* on 2L and *mScar* on 3R) enable simultaneous detection of LOH on different chromosome arms. (**d**) CRISPR targeting induces chromosome arm-specific LOH. sgRNA arrays targeting *Sfp24C1* (2L) or *CG5892* (3R) result in LOH exclusively on their respective chromosome arms. Points represent measurements in individual wing discs, columns represent the mean and error bars the standard deviation. (**e**) A system for high-throughput LOH screening *in vivo*. The *GMR11F02* enhancer allows highly selective expression of GFP in the pouch region of wing and haltere imaginal discs. (**f**) Control wing disc showing uniform GFP expression throughout wing pouch cells, with natural variations in expression level. (**g**) Example of LOH in a wing disc of a living larva. LOH events are readily detected as clearly defined patches lacking GFP fluorescence. Scale bar: 50 µm. (**h**) High-throughput screening for LOH on chromosome arm 2L reveals high on-target and undetectable off-target activity. 8197 wing discs expressing Cas12a^+^, GFP encoded on 2L and one of 525 HD12aCFD sgRNA arrays were screened for LOH. All 82 HD12aCFD lines targeting 2L between GFP and the centromere resulted in LOH, demonstrating robust on-target activity. Lines targeting 2L without LOH phenotype had target sites distal to GFP. Importantly, 443 HD12aCFD lines targeting other chromosome arms showed no reproducible LOH, indicating absence of detectable off-target activity on 2L (16% of the *Drosophila* genome). Sporadic low-level LOH events were not reproducible (Supplementary Data Fig. 2) and likely represent background rates.

We established high-throughput LOH screening using a tissue-specific enhancer to restrict GFP expression to *Drosophila* larval wing and haltere imaginal discs (35) (Fig. 2e and methods). This approach enables visualization of LOH in living animals with a conventional fluorescent stereoscope (Fig. 2f, g). We used this system to screen for LOH of a reporter inserted at cytogenetic map position 22C on chromosome 2. Induction of DSBs within the 21.4 Mb region (16% of the *Drosophila* genome) between this insertion site and the centromere (the screening interval) are expected to trigger LOH of the reporter. Our screen analyzed over 8,000 imaginal discs using 525 HD12aCFD lines that collectively encode 2,197 unique sgRNAs (Fig. 2h). Remarkably, all 82 sgRNA lines targeting genes located within the screening interval resulted in robust LOH (Fig. 2h). Moreover, of the 443 lines targeting genes outside the screening interval, 427 showed no LOH and 16 had LOH only in isolated wing discs (Fig. 2h, Supplementary Data Fig. 2). The latter could reflect either low-level off-target activity within the screening interval or stochastic background LOH unrelated to CRISPR. Off-target effects should be reproducible as they are caused by sgRNA homology, while stochastic LOH occurs randomly. To distinguish between these possibilities we rescreened these lines with larger sample sizes. This revealed no reproducible LOH patterns, indicating these rare events are unrelated to CRISPR activity (Supplementary Data Fig. 3). The frequency of such events is consistent with previous reports on stochastic LOH in *Drosophila* (36,37). To determine how LOH scores correlate with mutation frequencies, we sequenced target sites of sgRNAs that produced low but reproducible LOH signals. This revealed that the sensitivity of our LOH screening approach is comparable to or exceeding that of conventional Sanger sequencing of PCR amplicons (Supplementary Data Fig. 4). In summary, visualizing LOH enabled us to screen for Cas12a^+^ activity in combination with 525 sgRNA arrays across a 21.4 Mb interval of the *Drosophila* genome. This constituted to our knowledge the so far largest unbiased screen for nuclease activity *in vivo* and reveals both the robust activity and high specificity of multiplexed Cas12a mutagenesis.

### Multiplexed Cas12a^+^ gene targeting outperforms established Cas9-based methods

We then evaluated whether the HD12aCFD system enhances knock-out efficiency *in vivo* compared to the current state-of-the-art. To enable this comparison, we developed a toolkit comprising optimized, inducible *UAS-Cas12a^+^*transgenes and a whole-animal reporter system for nuclease activity mapping (Supplementary Data Figure 5). Using this toolkit, we performed both ubiquitous and tissue-specific CRISPR mutagenesis of endogenous genes, comparing HD12aCFD sgRNA arrays to established Cas9 sgRNA lines through quantification of loss-of-function phenotypes. First, we compared the effectiveness of both systems by targeting *white* (*w*) in neuronal photoreceptors. While Cas9-mediated mutagenesis produced highly variable mosaics with 50 ± 26% (mean ± standard deviation) of the tissue retaining *w* expression, gene targeting with HD12aCFD resulted in highly penetrant *w* disruption (96 ± 5%) (Fig. 3a, b). Next, we performed inducible mutagenesis of *Notch* and *neuralized* in adult intestinal stem cells. Both genes are known to control stem cell numbers by regulating cell proliferation (Fig. 3c). HD12aCFD-mediated targeting resulted in a more pronounced increase in dividing cells compared to Cas9 with sgRNAs pairs for both genes (Fig. 3d). Furthermore, targeting *smoothened* specifically in the pouch region of wing imaginal disc epithelia resulted in near complete loss of Smo protein with a HD12aCFD line, but a significant number of cells retained Smo in flies with two sgRNAs and Cas9 (Fig. 3e, f). These results demonstrate that gene targeting with Cas12a^+^ and multiplexed sgRNAs achieves superior gene disruption efficiency compared to established Cas9-based systems across diverse tissues. Additionally, Cas12a^+^ activity can be effectively controlled in time and space, as phenotypic changes remained confined to the targeted regions - a precision that has been challenging to achieve with Cas9 (4).

**Figure 3:**
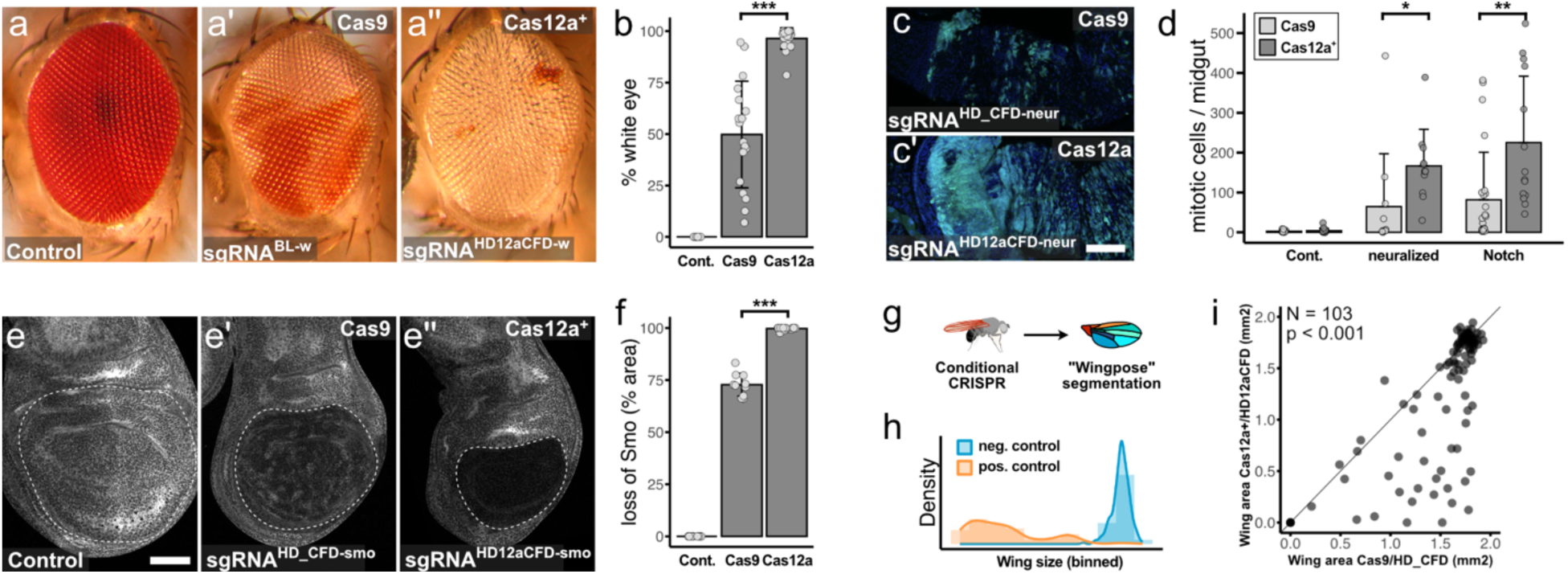
Cas12a^+^ and HD12aCFD arrays enhances gene knock-out efficiency compared to current Cas9-based systems. (**a-a’’**) The HD12aCFD system improves disruption of *white* (*w*) in the eye. Representative images of eyes from flies with no sgRNA (control) or Cas9 and a single sgRNA or Cas12a^+^ and a HD12aCFD array. All genotypes contain three functional copies of *w* due to transgene markers. (**b**) Quantification of white eye area. Individual data points show white area per eye, with mean (columns) and standard deviation (error bars). **p<0.001, independent two-sample t-test. (**c-c’**) Posterior *Drosophila* midgut showing stem and progenitor cells (ISCs/EBs, green) and DNA (blue). Targeting *neuralized* (*neur*) with Cas12a+/HD12aCFD increases ISC/EB numbers. Scale bar: 50 µm. (**d**) Quantification of mitotic cells (phospho-Histone3 positive) in the midgut. Loss of *neur* or *Notch* function leads to an increase in cell proliferation, with significantly stronger effects using Cas12^+^ and HD12aCFD sgRNA arrays. Points: mitotic cells per midgut; columns: mean per condition; error bars: standard deviation; **p<0.01, *p<0.05, independent two-sample t-test. (**e-e’’**) Immunofluorescent staining of Smo in wing imaginal discs. Conditional mutagenesis in the pouch region (dotted line) with Cas12a^+^ and HD12aCFD sgRNAs results in more effective gene knock-out. Note correlation between knock-out efficiency and pouch size. Scale bar: 50 µm. (**f**) Quantification of Smo protein loss. Data points show affected area per disc, with mean (columns) and standard deviation (error bars). ***p<0.001, independent two-sample t-test. (**g**) Schematic of the experimental system used in (h) and (i). Tissue-specific CRISPR results in wing specific mutagenesis. Wing size is analyzed with help of a custom Cellpose model (see methods). (**h**) Wing size distribution comparing Cas12a+ targeting of known wing morphogenesis genes (orange, n=33) versus control genes (blue, n=18). Clear phenotypic separation demonstrates specificity. See Supplementary Data Fig. 5 for details. (**i**) Comparative analysis of wing size phenotypes using Cas12a+/HD12aCFD versus Cas9/HD_CFD systems. Zero values indicate lethal phenotypes. Systems show significantly different effects on tissue size (Wilcoxon signed-rank test, V = 3915, n = 103, p < 0.001, two-tailed).

To more systematically assess HD12aCFD-mediated gene disruption efficiency, we used adult wing size as a quantitative phenotypic readout. We targeted 241 genes using conditional Cas12a^+^ and HD12aCFD sgRNA arrays, performed parallel imaging of multiple wings per genotype, and trained a widely-used cell segmentation algorithm (38) for automated segmentation of wings (Fig. 3g). Wing size remained largely unchanged when targeting negative control genes, but was strongly reduced upon disruption of known regulators of wing growth and morphogenesis (Fig. 3h, Supplementary Data Fig. 6). We compared our results to Cas9-mediated mutagenesis for the 103 genes with existing sgRNA lines in the HD_CFD library. HD12aCFD targeting produced significantly stronger phenotypes for the majority of wing size regulators (Fig. 3i). This included many genes known to be important for wing growth and morphogenesis, but also genes with previously unknown wing phenotypes (Supplementary Data Fig. 6). For example, mutagenesis of *trade embargo* (*trem*) with HD12aCFD sgRNA arrays, but not HD_CFD sgRNA pairs, resulted in a specific and highly penetrant loss of wing veins in the wing posterior (Supplementary Data Fig. 7). Perturbation of *trem* with additional sgRNAs and RNAi lines confirmed a role for *trem* in the wing, which had been missed in previous studies (39). Together, these results establish Cas12a^+^ with four sgRNAs as a superior system for generating loss-of-function alleles, achieving substantially higher knockout efficiency than established Cas9-based systems.

## Discussion

In this study, we developed an improved CRISPR gene ablation system. Our approach combines Cas12a^+^ and four arrayed sgRNAs targeting independent sites in each gene to overcome a general limitation of gene knockouts with CRISPR nucleases in multicellular systems - the generation of genetic mosaics that include cells with functional gene copies. Direct comparisons with current Cas9-based approaches demonstrate that this system not only increases knockout penetrance across diverse tissues, but also reveals phenotypes previously missed by conventional approaches. Importantly, systematic evaluation of potential limitations showed that the method maintains high specificity despite increased sgRNA multiplexing and is well tolerated *in vivo*, making it a robust platform for functional studies.

The superior performance of Cas12a^+^ with quadruple sgRNA arrays stems from both redundant and synergistic effects of multiplexed targeting. Redundancy of sgRNAs ensures robust editing - as demonstrated by 100% activity of all 82 on-target arrays in our LOH assay - in contrast to established single and double Cas9 sgRNA libraries that typically include 10 - 20% inactive lines (10–12). Additionally, multiplexed sgRNAs can act synergistically by creating deletions between target sites. While induction of such deletions require near-simultaneous DNA cutting and are rare with two sgRNAs, their frequency increases substantially when four sgRNAs targeting the same locus are used. Additionally, it is possible that differences in cleavage activity and kinetics between the used Cas9 and Cas12a variants influence overall performance (40).

Possible adverse effects of sgRNA multiplexing include toxicity from excessive DNA damage. Studies in mammalian cells have shown that cells with multiple target sites activate DNA damage checkpoints and are lost in pooled CRISPR screens (41,42). However, in these cases the gene copies are usually dispersed in the genome. We find that increasing sgRNA number from two to four at a single locus does not detectably enhance cell death. In contrast, targeting gene copies on multiple chromosomes, even with fewer total cut sites, significantly increases apoptosis and causes severe adult phenotypes. This suggests that the spatial distribution of cut sites, rather than their total number, is a major determinant of toxicity - possibly through the formation of chromosomal translocations when targeting multiple loci.

Multiplexing several sgRNAs could theoretically increase off-target effects. Because off-target activity varies widely between individual sgRNAs, meaningful characterization requires analysis of many guides. However, high-throughput assays to detect CRISPR activity at unknown genomic loci have been lacking. Our new method for visualizing LOH in living animals fills this technical gap. Since LOH is a widespread repair outcome of DSB repair in diploid organisms (29–31), this assay should be broadly applicable to other systems. Screening more than 2,000 sgRNAs in 525 arrays across 21 megabases of the *Drosophila* genome failed to detect any off-target activity. This confirms that the high specificity of Cas12a observed *in vitro* (43) is maintained during *in vivo* multiplexed editing.

*Drosophila* was the first organism in which targeted gene editing was achieved (44) and has proven to be an excellent model for genome engineering approaches that later proved valuable in other animals (4,45). This broad transferability stems from the fact that Cas proteins function effectively across different organisms, and that the mechanisms of DSB repair are highly conserved during evolution. The improved efficiency and high specificity of multiplexed Cas12a gene targeting therefore has implications beyond basic research applications. For example, current gene drive systems, which hold promise for controlling vector-borne diseases, are limited by the emergence of resistance alleles. Recent studies have demonstrated that sgRNA multiplexing can substantially delay resistance (46,47) suggesting that our system could accelerate the development of more robust genetic tools for disease control. Ultimately, this work not only provides a blueprint for developing enhanced gene knockout systems across multicellular organisms, but also contributes to a broader toolkit for addressing both fundamental questions in biological research and pressing societal needs.

## Material and Methods

### Plasmids

#### Plasmid construction

Plasmid cloning was performed using PCR amplicons generated with Q5 PCR polymerase (New England Biolabs). DNA fragments were assembled by either In-Fusion cloning (Takara) or by GoldenGate cloning using BbsI-HF and T4 DNA Ligase (both New England Biolab) using standard procedures or by extension overlap PCR (see below). PCR conditions followed manufacturer’s recommendations. Newly cloned inserts were confirmed by Sanger sequencing. Sequences of plasmids and primers can be found in Supplementary Table 1. Plasmids are available from the European Plasmid Repository (https://www.plasmids.eu) and Addgene (https://www.addgene.org).

##### pSTAR(1-6x)

The reporter plasmids for multiplexed cutting (<u>s</u>gRNA target site <u>ar</u>ray (STAR)) were generated by cloning a fragment of the *lacZ* coding sequence into a plasmid backbone containing *AmpR* and *mini-white* selection genes, as well as *attB* and *ori*, that was derived by BamHI digestion of plasmid *pCFD6* (4). The *lacZ* fragment lacks both a promoter and start codon and is presumed to be transcriptionally inactive. Afterwards, the sequence TTTCCAGGGAACTCCCATCCACCATGGCTGGTATTCC, which can be targeted by a Cas12a sgRNA (CAGGGAACTCCCATCCACCATGG) or Cas9 sgRNA (GAACTCCCATCCACCATGGC), was introduced by overlap extension PCR. Two PCR amplicons were generated with an overlapping sequence containing the sgRNA target site (see Supplementary Table 1 for primer sequences). Equimolar amounts of the two PCR amplicons were mixed and subjected to PCR for 15 cycles. Then, primers annealing to the end of the fused PCR amplicon were added and full length products amplified for another 20 PCR cycles. PCR amplicons were separated on a 1% agarose gel, full length amplicons were excised, purified and cloned into BamHI digested pCFD6 by In-Fusion cloning.

##### act5C-mScarlet

A double-stranded DNA fragment encoding the *act5C* promoter, *mScarlet* coding sequence and *act5C* 3’UTR was synthesized from Integrated DNA Technologies. It was assembled with a plasmid backbone fragment containing *AmpR* and *mini-white* selection genes, as well as *attB* and *ori*, that was obtained by BamHI digestion of plasmid *pCFD6* (4) by In-Fusion cloning. The plasmid is available from the European Plasmid Repository (EPR#897) and Addgene (Plasmid 230906).

##### GMR11F02-Gal4 UAS-GFP

A fragment encoding the *GMR11F02* regulatory element and Gal4 coding sequence was PCR amplified with primers *GMR11F02G4fwd* and *GMR11F02G4rev* from plasmid *pGMR11F01-Gal4* (a gift from Ryan Loker and Richard Mann, Columbia University, New York). The fragment was then inserted into *pUAST-EGFP* digested with XhoI by In-Fusion cloning. The plasmid is available from the European Plasmid Repository (EPR#898) and Addgene (Plasmid 230907).

##### UAST-u^M^Cas12a^+^nls^1x^

Plasmid *UAST-u^M^Cas12a^+^nls^1x^* was derived from *UAS-LbCas12a* (25). An upstream open reading frame was PCR amplified from *UAS-u^M^Cas9* (10) with primers *uMCas12afwd* and *uMCas12arev* and gel purified and cloned into *UAS-LbCas12a* digested with EcoRI by In-Fusion cloning.

##### UAST-FRT-GFP-FRT-u^M^Cas12a^+^nls^1x^

Plasmid *UAST-FRT-GFP-FRT-u^M^Cas12a^+^nls^1x^*was derived from *UAST-u^M^Cas12a^+^nls^1x^*. The plasmid was digested with EcoRI-HF and HindIII and the sequence encoding the FRT-GFP-FRT cassette was synthesized by Integrated DNA Technologies and inserted by In-Fusion cloning. The plasmid is available from the European Plasmid Repository (EPR#899) and Addgene (Plasmid 230908).

##### UAST-u^M^Cas12a^+^nls^2x^

Plasmid *UAST-u^M^Cas12a^+^nls^2x^* was derived from *UAST-u^M^Cas12a^+^nls^1x^*. The sequence encoding an additional C-terminal nucleoplasmin nuclear localization signal was ordered as double-stranded DNA from Integrated DNA Technologies and inserted into *UAST-u^M^Cas12a^+^nls^2x^* digested with NdeI and HpaI by In-Fusion cloning. The plasmid is available from the European Plasmid Repository (EPR#900) and Addgene (Plasmid 230909).

##### pGRACE

Plasmid *pGRACE* was derived from pAct5c-LbCas12a^+^ (25). The backbone was linearized with EcoRI and XhoI. The sequence encoding the uORF with binding sites for Cas12a and Cas9 sgRNA flanked by microhomology and the fluorescent protein was synthesized by Integrated DNA Technologies and inserted by In-Fusion cloning. The plasmid is available from the European Plasmid Repository (EPR#901) and Addgene (Plasmid 230910).

#### Cloning of sgRNA plasmids

Individual sgRNA plasmids were constructed as previously described (25). The spacer sequences of sgRNAs are listed in Supplementary Table 1.

### sgRNA library design

All guides used in this study have been designed using the CRISPR Library Designer (v. 1.5) (48). Guides were designed against the *D. melanogaster* genome release BDGP6.22, allowing only guides in coding sequences with less than 10 predicted off-targets of an edit distance larger than 2. For aligning crRNA target sites back to the reference genome bowtie was used in very-sensitive mode and the first 3 PAM distal bases were ignored. The PAM requirement was set to a 5’ adjacent TTTN. As basis for design served a list of ENSEMBL identifiers covering all protein coding genes in BDGP6.22. Raw, designed crRNA targets were then further processed in a custom R pipeline to yield optimized crRNA target pairs for combinatorial mutagenesis as follows.

The CLD output files were filtered according to the design criteria. In the following steps, relevant columns from the large CLD output table were selected. Everything irrelevant to the list of target genes was removed, and the ‘Number of Hits’ column was adjusted to represent off-target count by subtracting one count. After that, all sequences containing BbsI (GAAGAC) and BsaI (GGTCTC) restriction sites, non-unique crRNA targets, and crRNA targets adjacent to a TTTT PAM were removed. Additional selection criteria for the remaining pool of sgRNA designs were annotated as follows: the number of targeted protein-coding transcripts, whether the sgRNAs target the first half of the target gene, and the GC content of sgRNA sequences. Only those sgRNAs that hit at least max(#transcripts)-1 transcripts were kept, and the average gene expression for each transcript hit by a putative crRNA was annotated. Designs that unintentionally hit a coding sequence of a different gene than they were designed for were filtered. Further, small genes that are contained in overarching giant genes like *kirre*, *Raf*, or others were excluded. Each sgRNA was annotated on how many CDS it targets and if it targets in the first half of the target gene. To check the pool for sanity, the sgRNAs were written in GFF3 format. This file could then be loaded into an arbitrary genome browser. Next, 180 random pairs of sgRNAs were drawn for each gene and scored by optimizing pair distance, microhomology score (49), exon expression, isoform coverage, off-target counts, GC content, and exon position relative to the transcript. The two top-scored crRNA pairs were chosen for further quality controls by manual and automatic inspection.

### Cloning of the HD12aCFD sgRNA library

Cas12a sgRNAs were cloned into the BbsI site of the expression vector *pCFD8*, which has been previously described (European Plasmid Repository EPR#32; Addgene #140619)(25). Oligonucleotides encoding four sgRNAs, as well as primer binding and BbsI restriction sites at either end were synthesized by Twist Bioscience. The oligonucleotide design can be found in Supplementary Table 1. Oligos were resuspended in dH_2_O and amplified by PCR using primers *Cas12arrayAMPfwd2* and *Cas12arrayAMPrev2*. PCR products were separated on a 1% agarose gel, excised, purified and cloned into *pCFD8* by GoldenGate cloning using standard conditions. DNA was transformed into chemically competent E. coli (Stellar Silver competent cells, Takara) and plated on LB agar plates containing Carbicilin. After overnight incubation at 37C, ten individual colonies were picked for validation purposes and all other colonies were pooled. DNA was extracted using QIAprep Spin Miniprep Kit (Qiagen) following manufacturer’s instructions. DNA from individual colonies was validated by Sanger sequencing. When at least 9 of the 10 tested colonies contained unique, correct sgRNA plasmids we proceeded with injection of the DNA from the pooled colonies into *nos-PhiC31; attP40* transgenic *Drosophila* embryos.

### Visualization of CRISPR-edited PCR amplicons

To visualize diversification of the target locus by CRISPR editing, reflecting the heterogeneity of the edited sequences, genomic DNA was extracted from individual flies as previously described (25). The target locus was amplified by PCR using locus-specific primers (Supplementary Table 1) with Q5 High-Fidelity DNA Polymerase (NEw England Biolabs) and no more than 30 PCR cycles under standard conditions. Amplicons were separated on 1% agarose gels. Note that amplicon pools containing diverse CRISPR edits typically form heteroduplexes during PCR, which results in the formation of a smear towards higher- and lower molecular weight products on the gel.

### Identification of CRISPR alleles by Sanger sequencing

To identify CRISPR-induced alleles genomic DNA was extracted from individual flies as previously described (25) and the target locus was amplified by PCR using locus-specific primers (Supplementary Table 1) that annealed typically 300-500 bp 5’ and 3’ of the sgRNA target site. PCR amplicons were purified using paramagnetic beads (50) and sent for Sanger sequencing (Eurofins Genomics).

### Fly stocks and husbandry

Transgenic *Drosophila* strains used or generated in this study are listed in Supplementary Table 2. *Drosophila* were cultured on standard cornmeal food media in an incubator set to 27°C with a 12h light - 12h dark cycle and 60% relative humidity. Adult flies were transferred to fresh food vials every 3-4 days.

### Transgenesis

Transgenesis was performed with the PhiC31/attP/attB system and plasmids were inserted in landing sites on the second or third chromosome as indicated for each line. Microinjection of plasmids into *Drosophila* embryos was carried out using standard procedures. Transgenesis of crRNA plasmids was typically performed by a pooled injection protocol, as previously described (25). Briefly, individual plasmids were pooled at equimolar ratio and DNA concentration was adjusted to 250 ng/μL in dH_2_O. Plasmid pools were microinjected into blastoderm embryos, raised to adulthood and individual flies crossed to P{ry[+t7.2] = hsFLP}1, y[1] w[1118]; Sp/CyO-GFP or P{ry[+t7.2] = hsFLP}1, y[1] w[1118];; Sb/TM6B. Transgenic offspring were identified by eye color and individual transgenic flies from pooled plasmid injections were genotyped as previously described (25). Flies were crossed to balancer flies and stable transgenic stocks were generated.

### Immunofluorescence staining

*Drosophila* larvae were dissected in ice-cold PBS and fixed in 4% paraformaldehyde for 25 minutes at room temperature. Tissue was washed with PBT (PBS with 0.3% Triton-X100) and blocked with 1% heat-inactivated normal goat serum in PBT. Antibodies were diluted in PBT and tissue was incubated overnight at 4°C. Antibodies used were anti-cleaved DCP1 (diluted 1:600, Cell Signaling), anti-phosphorylated Histone 3 (Cell Signaling) and anti-Smoothened (DSHB, 1:200). After serial washes in PBT tissue was incubated in a secondary antibody (labeled with Alexa Fluorophores, diluted 1:600, Life Technologies) and Hoechst (diluted 1:2000) in PBT for 2 hours at room temperature. Following serial washes tissue was mounted in Vectashield (Biozol Diagnostica).

### Imaging

Tissue was imaged on a Leica SP8 laser-scanning confocal microscope with an oil 40x/NA1.4 lens using the sequential scanning mode. Image processing and analysis was performed with FIJI. Imaging of adult flies or wings was performed with a Leica M165 FC stereomicroscope equipped with a Leica DFC295 camera.

### Loss of heterozygosity screening

LOH was monitored using heterozygous transgenes encoding fluorescent proteins. Upon LOH of the chromosome arm on which the transgene is located, cells gain or lose fluorescence, which can be visualized using a fluorescent microscope or stereo microscope. Plasmids were inserted at the distal end of the chromosome, either at cytogenetic position 22C on chromosome 2 or at 99F on chromosome 3. Animals were also transgenic for *act5C-Cas12a^+^* and HD12aCFD sgRNAs, and LOH was assessed in wing imaginal discs at larval 3rd instar wandering stage. We confirmed that LOH is also readily detected in other tissues, such as the brain and the intestine.

To assay LOH with high resolution the *act5C-mScarlet* plasmid was used, which results in ubiquitous and homogenous expression of mScarlet. Third instar larvae were dissected on ice, and wing discs were fixed in 4% paraformaldehyde in PBS at room temperature for 25 minutes. Nuclei were stained with 1 µg/ml Hoechst for 20 minutes and discs were mounted in Vectashield mounting medium. Imaginal discs were imaged with a Leica SP8 laser-scanning confocal microscope with an oil 40x/NA1.4 lens. Images were analyzed in Fiji by outlining all areas that lost fluorescence (this measure being less ambiguous than gain of fluorescence, which can be obscured by natural variation in fluorescence, e.g. in tissue folds) and using the measure function to quantify their size. Note that the area that cells with LOH occupy in the tissue reflects both the frequency of LOH (number of cells that gain or lose reporter expression) and the time point of occurrence (number of cell divisions after LOH). While the *act5C-mScarlet* reporter is ideal to observe LOH in isolated tissues by high resolution imaging, the ubiquitous reporter cannot reliably detect LOH in whole animals, as the multilayered topology obscures LOH events in individual tissues.

For high-throughput LOH screening in living animals the *GMR11F02-Gal4 UAS-GFP* reporter was used, which is specifically expressed in the pouch region of the wing and haltere imaginal discs. Larvae were immersed in a drop of tap water and observed under a Leica M165 FC fluorescent stereo microscope with 6.3x magnification. For consistency all screening was performed by the same observer (F.P.), who was blinded to larval genotypes. Typically 10 to 20 larvae were scored per genotype. The GFP signal from the *GMR11F02-Gal4 UAS-GFP* reporter in the disc pouch shows natural variation, with stronger signal typically observed at the dorsal edge of the expression domain and weaker fluorescence along the dorsal-ventral boundary. Therefore, the observer focused on larger patches of cells that lost fluorescence. The observer scored the level of LOH per wing disc on a scale from 0 (no LOH) to 3 (strong LOH).

### Drosophila wing size phenotyping

Dissected *Drosophila* wings were mounted on glass slides and imaged using a Leica M165 FC stereo microscope. Images were used to retrain Cellpose’s cyto2 model (38) using human-in-the-loop capabilities and parameters: diameter 140, flow 1, and cell prob. 1. Using this model (‘fly_wings’) wings were segmented, outlines were exported to Fiji and wing size was measured using the Measure feature.

### Data analysis and visualization

Data analysis was performed in R using the Tidyverse package. Graphs were created using ggplot2. Microscope images were prepared using Fiji. Figures were created using Affinity Designer.

## Contributions

F.P and M.B. conceived the study. F.P., M.A.B., J.Z. designed, performed and analyzed experiments. M.S., A.V.B., A.C.M., E.R., A.P., L.G., J.H., L.B.M.K., B.W., M.H. performed experiments. F.H. designed the sgRNA library and trained the Cellpose model. M.B. acquired funding. F.P. and M.B. supervised the work. F.P. wrote the paper. All authors reviewed and edited the draft.

## Acknowledgements

We would like to thank Ryan Loker and Richard Mann (Columbia University, USA) for plasmids and Erich Brunner (University of Zürich, Switzerland) for advice about construction of the GRACE reporter. Roman Doll (University of Oxford, UK) for discussions and comments on the manuscript. Jianing Zhang, Nikola Knoll, Claudia Strein and Alma Spahic for discussions and support. The High-Throughput Sequencing Unit of the Genomics and Proteomics Core Facility and Light Microscopy Facility at DKFZ for support. This work has in part been supported by grants from the European Research Council (DECODE) and the German Research Foundation (SFB1324).

## Supplementary Data Figures

**Supplementary Data Figure 1:**
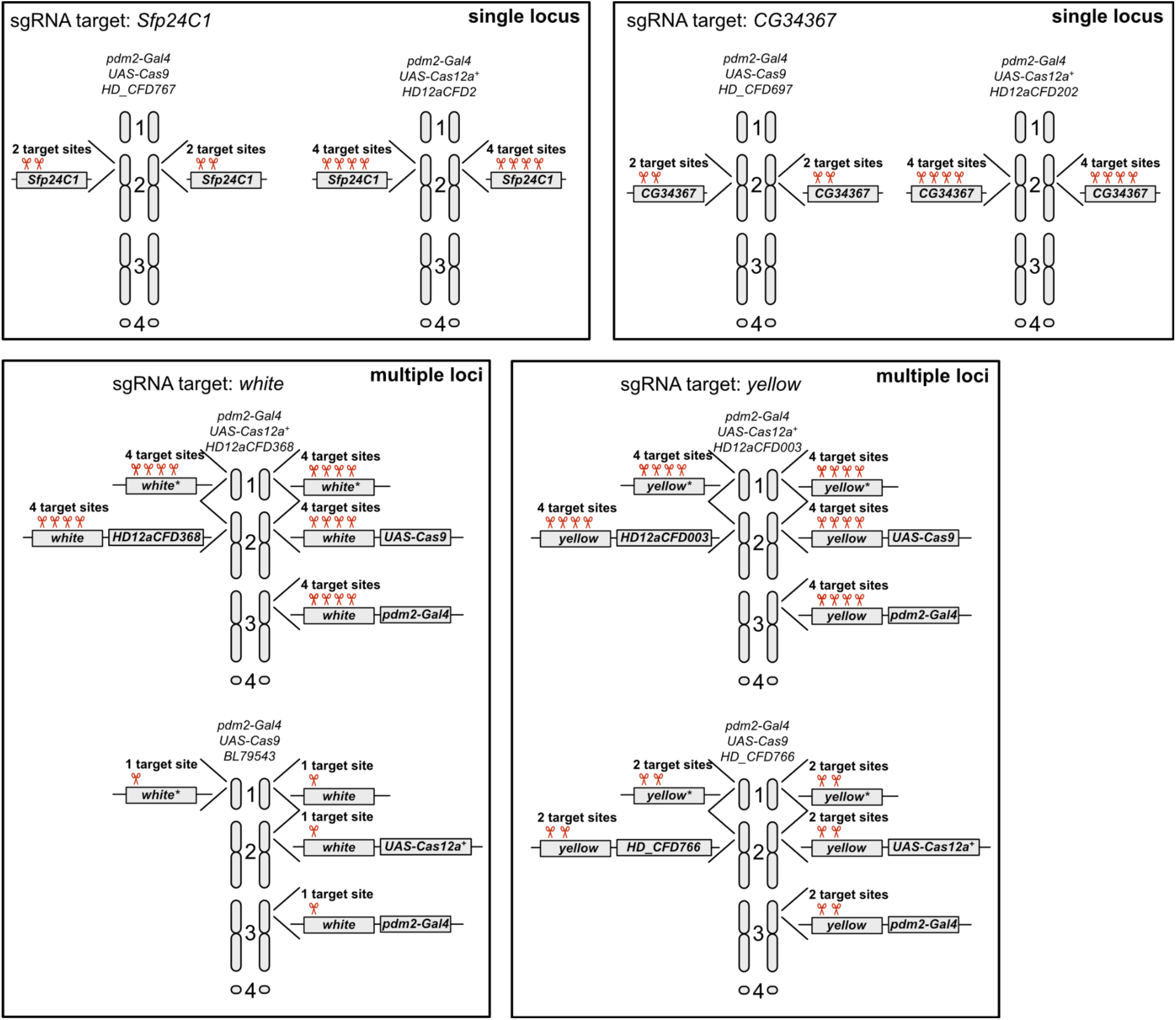
Genotypes and sgRNA target sites of strains used to test for DSB induced toxicity. Schematic of the genotypes of each strain used in Figure 1 f and g. sgRNA target sites are indicated by red scissors. Target genes *Sfp24C1* and *CG34367* are both present in a single copy on chromosome arm 2L. They were chosen because they are assumed to be non-essential for cells of the wing epithelium and LOH experiments showed that they are highly active. The genes *white* and *yellow* are used to mark transgenes or the landing sites they are inserted in and are therefore present on multiple chromosomes. Note that to target *white* with Cas9 we used a line encoding a single sgRNA (BL79543, a *white* targeting HD_CFD line does not exist). This transgene is marked by *vermillion* instead of *white*. Despite the low number of target sites this line still mediates increased cell death in the wing imaginal disc (Fig. 1g).

**Supplementary Data Figure 2:**
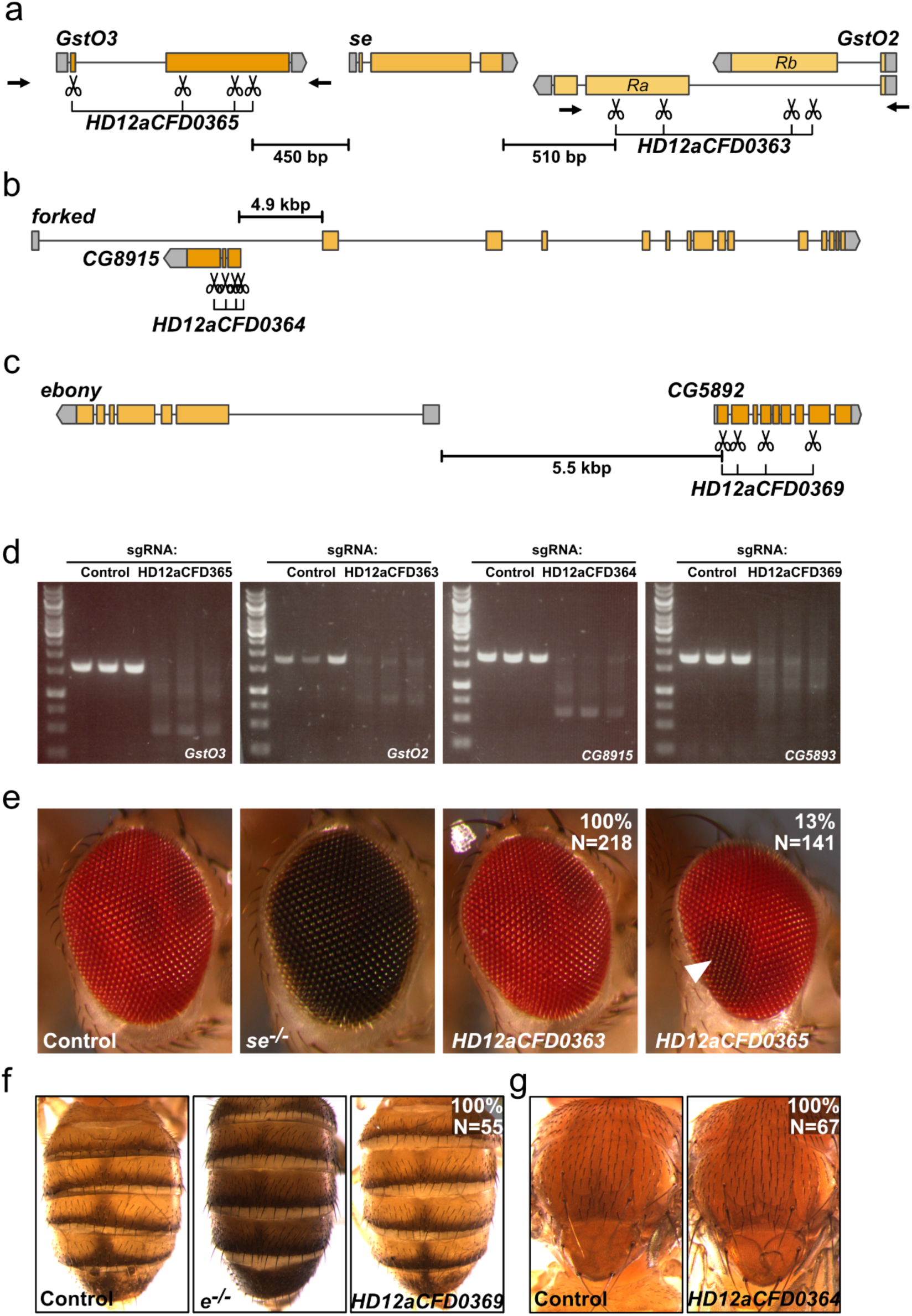
Deletions extending beyond the target locus are rare events. **(a-c)** Schematic of the genomic regions showing the gene with visible loss-of-function phenotype in the middle and the neighboring genes targeted by each sgRNA array. **(a)** The *sepia* (*se*) gene is shown with adjacent *GstO3* and *GstO2* loci. sgRNA arrays *HD12aCFD0363* and *HD12aCFD0365* target sequences in close proximity to *se*, with the closest sgRNA in *HD12aCFD0365* positioned just 450 bp upstream and *HD12aCFD0363* 510 bp downstream of *se*. Scissors symbols represent individual sgRNA target sites. Black arrows indicate PCR primer positions. **(b)** Genomic organization of the *forked* (*f*) locus and neighboring *CG8915* gene, with *HD12aCFD0364* target sites located 4.9 kb away. **(c)** Arrangement of the *ebony* (*e*) gene and *CG5892*, separated by 5.5 kb, with *HD12aCFD0369* target sites indicated. **(d)** CRISPR editing outcomes visualized by agarose gel electrophoresis. PCR amplification of target regions from flies expressing Cas12a^+^ and the indicated sgRNA arrays shows substantial diversification compared to control samples (expressing unrelated sgRNA). Each condition is shown in triplicate, revealing edited amplicons of varying sizes, with few maintaining the original wild-type length. **(e)** Eye phenotypes resulting from CRISPR targeting near the *se* locus. Control flies show normal red eye coloration (N>100), while *se^−/−^* mutants display characteristic dark eye pigmentation (N>100). *act5C-Cas12a^+^ HD12aCFD0363* flies (N=218) show no disruption of *se* function, while 13% of flies expressing *HD12aCFD0365* (N=141) exhibit small patches of dark eye tissue (white arrowhead), consistent with mosaic *se* disruption. This correlates with *HD12aCFD0365* proximity to *se*’s transcriptional start site and potential regulatory elements. **(f)** Abdominal pigmentation phenotypes showing normal patterning in control flies, darker pigmentation in *e^−/−^* mutants, and wild-type appearance in *act5C-Cas12a+ HD12aCFD0369* flies, indicating no disruption of *e* function despite nearby targeting. **(g)** Thoracic bristle morphology in control and *act5C-Cas12a+ HD12aCFD0364* flies, showing no evidence of *f* gene disruption despite targeting of adjacent sequences.

**Supplementary Data Figure 3:**
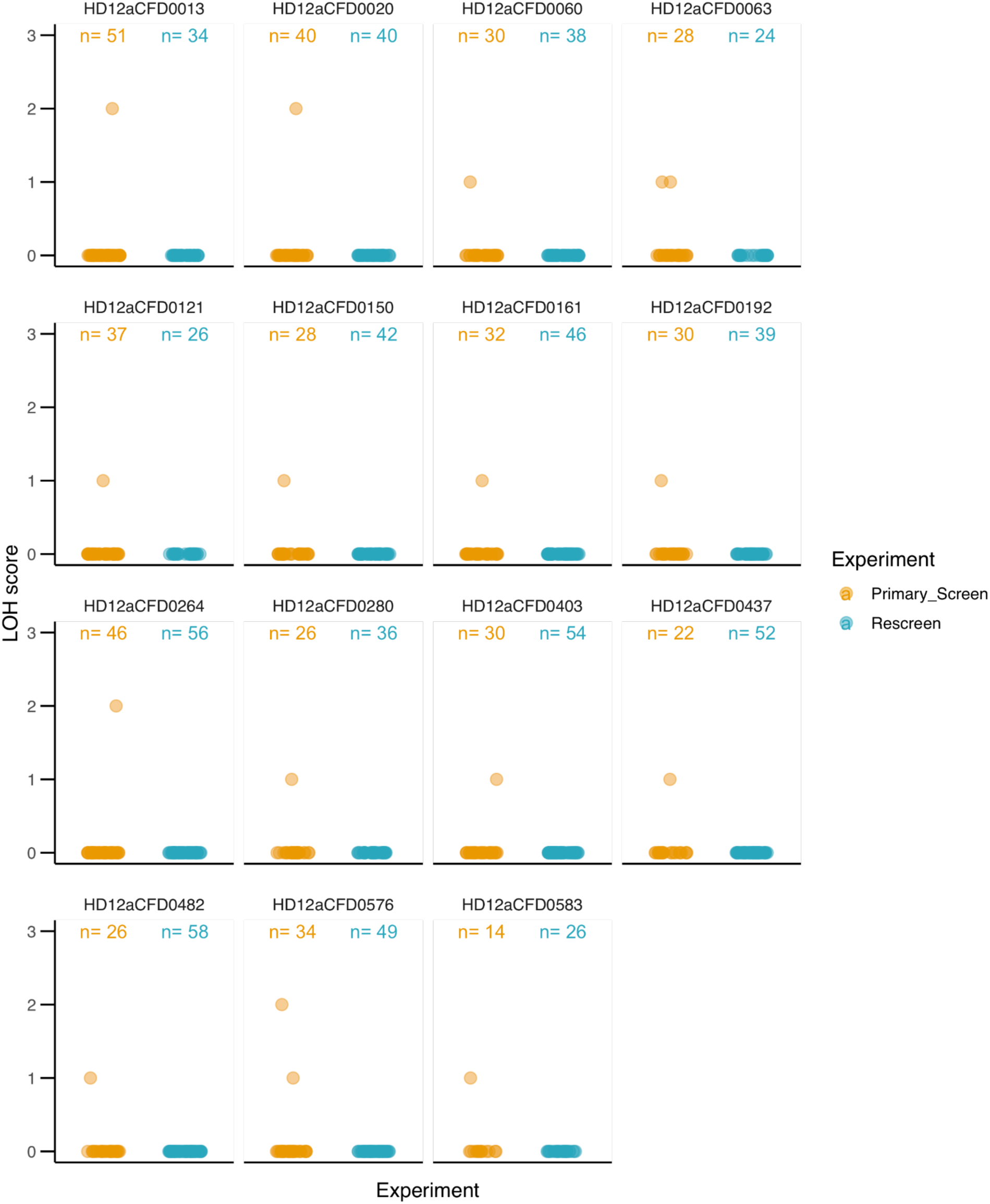
Sporadic LOH is not reproducible. HD12aCFD lines that resulted in sporadic LOH in the primary screen (presented in Fig. 2h) were rescreened using the same conditions. No LOH was detected, suggesting that the initial signal does not represent CRISPR off-targets, but likely either stochastic LOH or experimental error during the high-throughput screen.

**Supplementary Data Figure 4:**
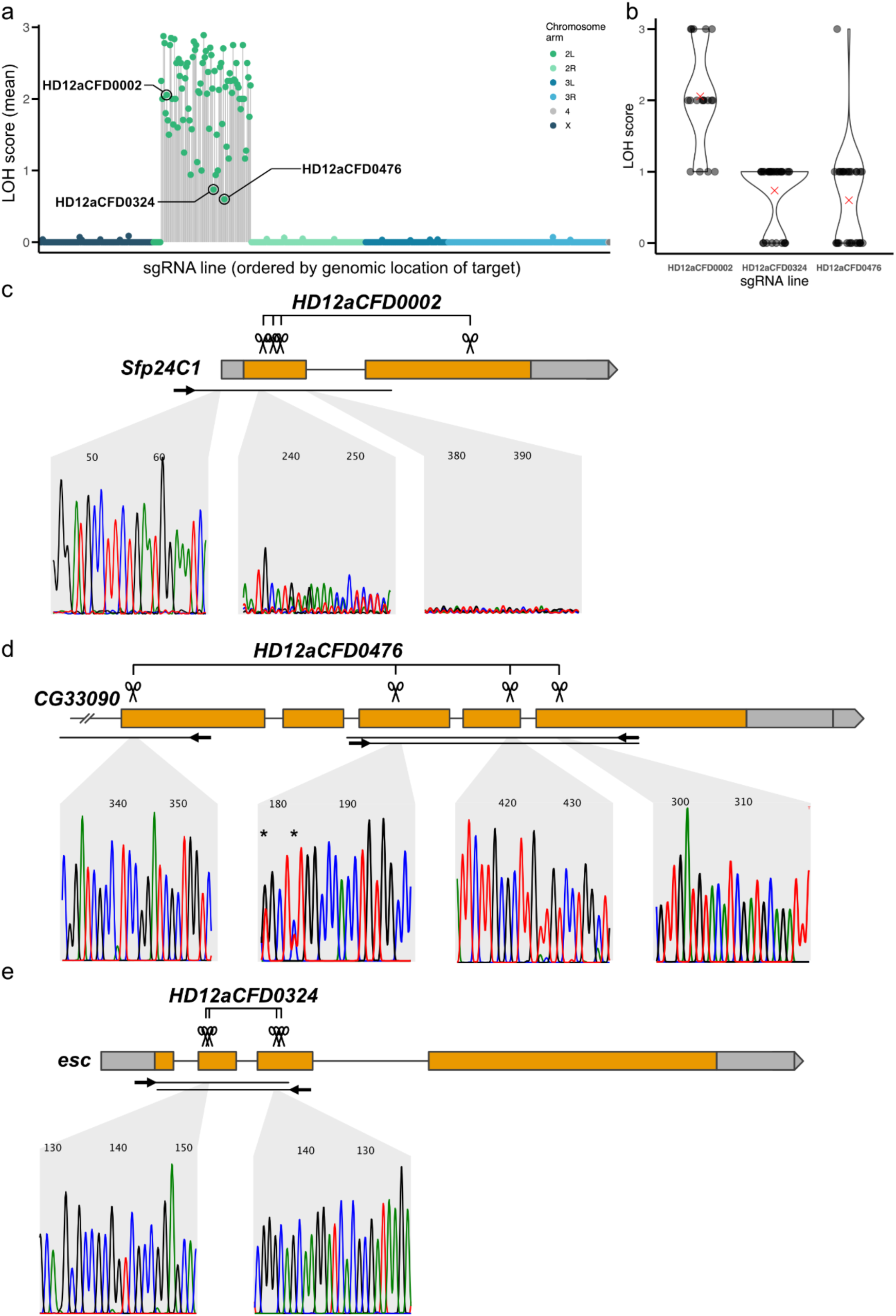
sgRNA lines that mediate low on-target mutagenesis result in reproducible LOH. (**a**) Distribution of mean LOH scores across all tested sgRNA lines, ordered by genomic location of their targets (Figure 2h). Highlighted are the three sgRNA lines (HD12aCFD0002, HD12aCFD0324, and HD12aCFD0476) selected for detailed analysis. (**b**) Loss of heterozygosity (LOH) scores for selected sgRNA lines. The violin plots show the probability density of the data at different values. Individual data points corresponding to individual wing imaginal discs (N = 18 - 34 per line) are shown as black dots. Red crosses indicate the mean LOH score for each sgRNA line. (**c** - **e**) Genomic context and sequencing traces for the three selected lines. Gene models show exons (orange boxes), untranslated regions (gray boxes), and sgRNA target sites (scissors symbols). Representative Sanger sequencing chromatograms are shown below each gene model. HD12aCFD0002 targeting *Sfp24C1* results in robust LOH (a) and strong sequence diversification at and behind the sgRNA target sites (c). HD12aCFD0476 targeting *CG33090* (d), and HD12aCFD0324 targeting *esc* (e) result in low, but reproducible LOH and show minimal to no detectable sequence diversification at the sgRNA target sites. Asterisks in panel (d) indicate sequence polymorphisms that are also found in a control genotype.

**Supplementary Data Figure 5:**
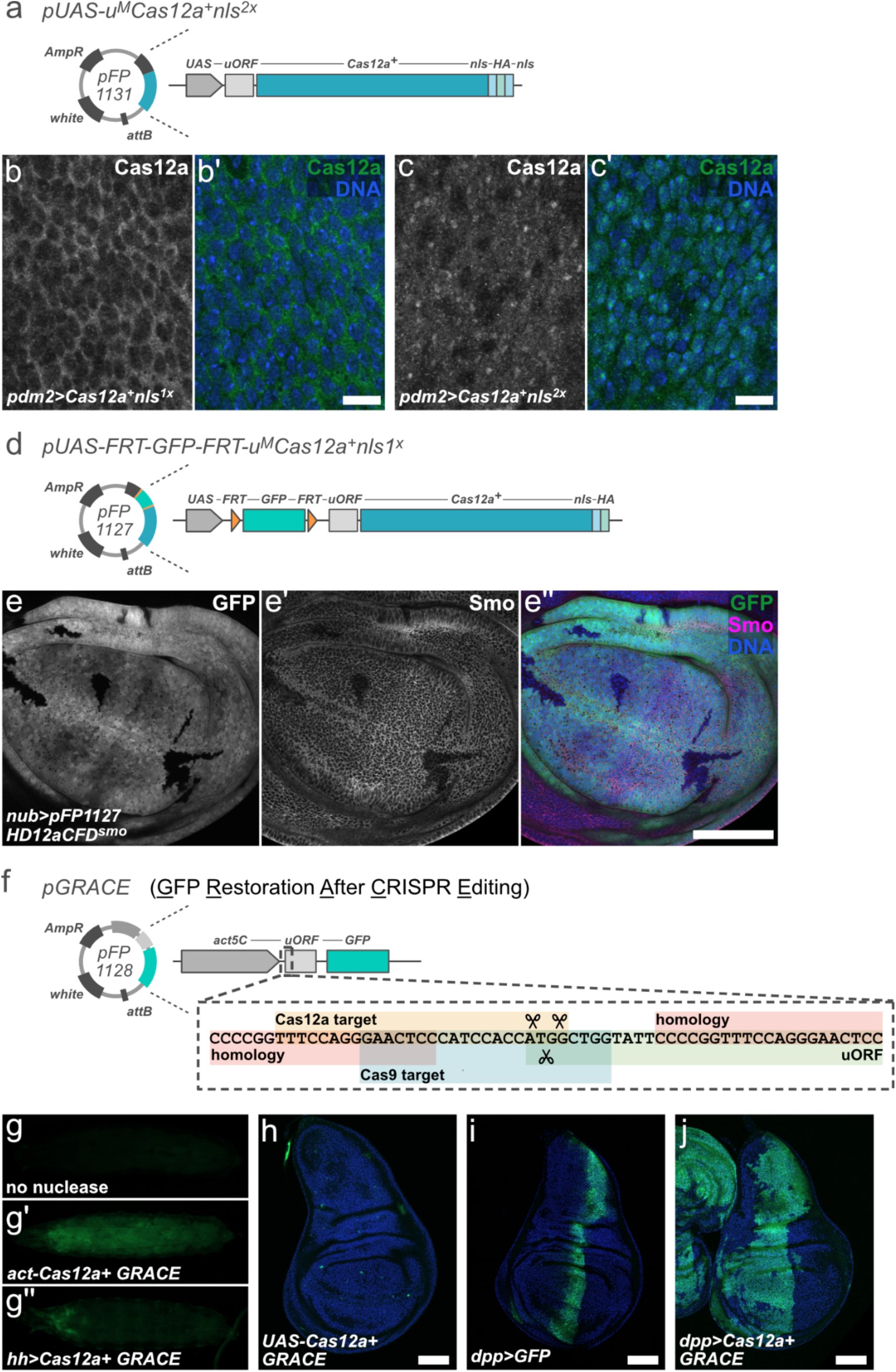
Transgenic tools for conditional Cas12a^+^ mutagenesis. (**a**) Schematic representation of the *pUAS-u^M^Cas12a^+^nls^2x^*construct (pFP1131) containing UAS-driven Cas12a with two nuclear localization signals (nls). The upstream open reading frame (uORF) attenuates Cas12a^+^ expression (10) and additional NLSs have been shown to increase editing efficiency in mammalian cells (51,52). (**b - b’**, **c - c’**) Immunofluorescence showing Cas12a expression (grayscale and green) with DNA counterstain (blue) in wing discs. Scale bars: 10 μm. A previously described Cas12a^+^ construct (25) with a single NLS localizes predominantly to the cytoplasm (b). Cas12a^+^ encoded on *pFP1131* localizes more efficiently to the nucleus (c). (**d**) Schematic of the *pUAS-FRT-GFP-FRT-u^M^Cas12a^+^nls^1x^* construct (pFP1127) featuring a FRT-flanked GFP transgene upstream of Cas12a for inducible expression through FLP expression. (**e - e’’**) Cas12a^+^ editing with spatial and temporal control. Immunofluorescence showing GFP (grayscale/green), Smoothened (Smo) (grayscale/magenta), and DNA (blue) *in hs-FLP nub-Gal4 UAS-pUAS-FRT-GFP-FRT-u^M^Cas12a^+^nls^1x^ HD12aCFD^smo^*wing imaginal discs approximately 60 h after a limited heat shock. Smo expression is selectively and efficiently lost in cells that excised the FRT-GFP cassette. Scale bar: 50 μm. (**f**) *pGRACE* (pFP1128) is a reporter to visualize gene editing activity *in vivo*. It encodes a ubiquitous *act5C* promoter and a GFP open reading frame separated by an uORF, which suppresses GFP translation. Upon CRISPR gene editing the start codon of the uORF is removed and GFP expression is derepressed. Detailed sequence of the Cas12a/Cas9 sgRNA target regions and homology regions to bias repair is shown in the expanded view. (**g - g’’**) GFP expression in larva without nuclease (g), with *act5C-Cas12a^+^ GRACE* (g’), and *hh-Gal4 UAS-u^M^Cas12a^+^nls^2x^ GRACE* (g”). All genotypes also express the Cas12a GRACE-sgRNA. GRACE readily reveals ubiquitous Cas12a activity when expressed from the *act5C* promoter and tissue-specific editing when expression is driven by *hh-Gal4*. (**h** - **j**) Wing disc GFP expression patterns showing *UAS-u^M^Cas12a^+^nls^2x^ GRACE* (h), *dpp-Gal4 UAS-GFP* (i), and *dpp-Gal4 UAS-u^M^Cas12a^+^nls^2x^ GRACE* (j). In the absence of a Gal4 driver only very few cells express GFP, presumably reflecting low levels of leaky Cas12a+ expression (h). Acute expression of GFP from a *UAS-GFP* transgene driven by *ddp-Gal4* is visible in a domain along the anterior-posterior boundary (i). In contrast, GRACE reveals that gene editing is active in a much broader domain comprising most of the anterior compartment (j). This is likely due to broader expression of *dpp-Gal4* in early development. Scale bars: 50 μm.

**Supplementary Data Figure 6:**
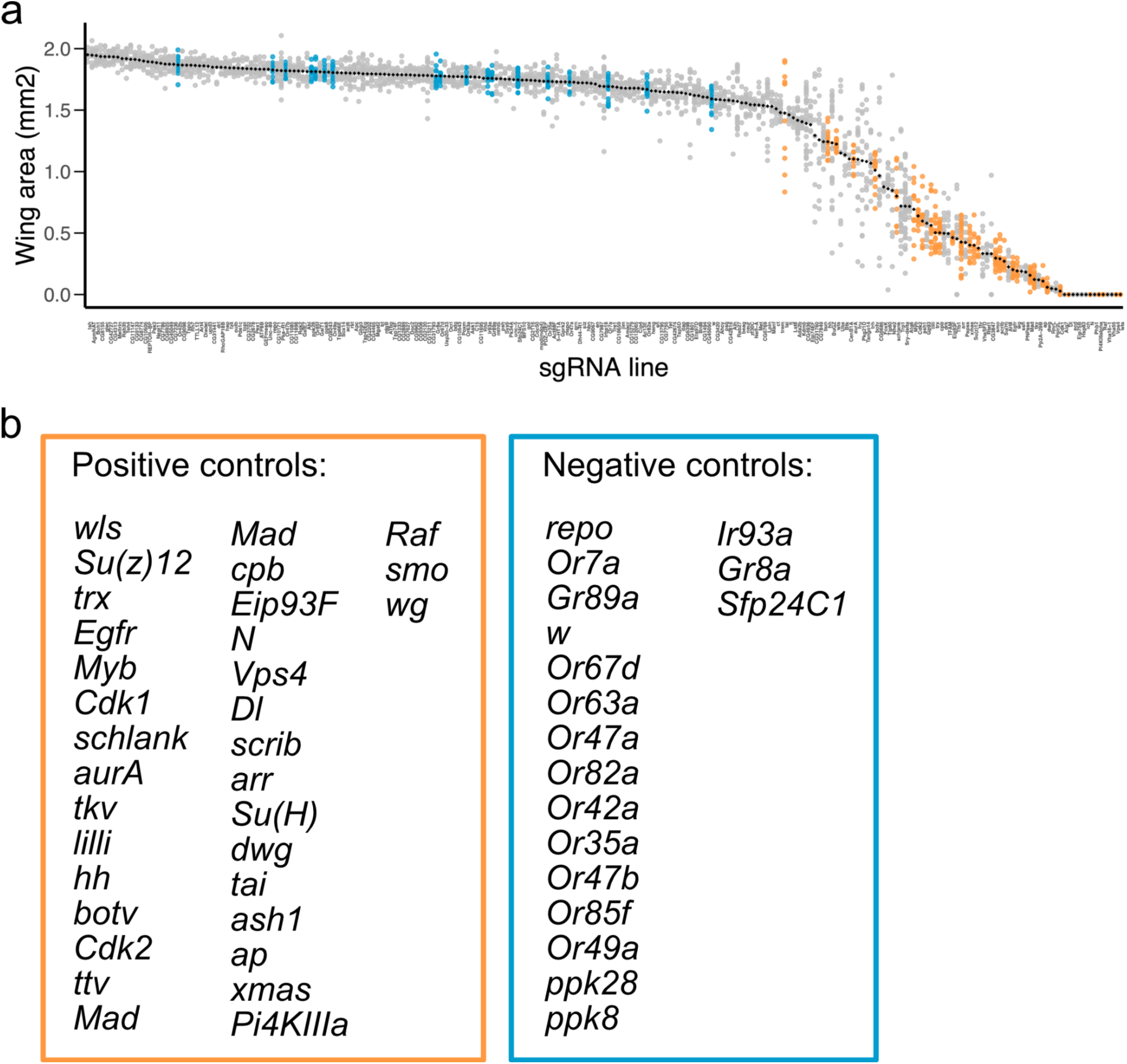
Tissue-specific CRISPR screening in wings with Cas12a+ and HD12aCFD sgRNA arrays. (**a**) Size of individual wings from animals expressing *pdm2-Gal4 UAS-u^M^Cas12a^+^* and *HD12aCFD* sgRNA arrays targeting the indicated genes. Each dot presents an individual wing (with typically between 10 - 20 wings measured per genotype) and the black rectangle represents the mean. Selected lines targeting genes known to affect or not affect wing morphogenesis are colored in orange or blue according to the color scheme in (b). While sgRNA targeting negative control genes have minimal effects on wing size, sgRNAs targeting positive control lead to a clear reduction in wing size. Phenotypes that were lethal are classified as zero and visible towards the right of the graph. (**b**) List of genes considered as positive (orange rectangle) or negative (blue rectangle) controls based on previous literature. Wing size of these lines is also plotted in Figure 3h.

**Supplementary Data Figure 7:**
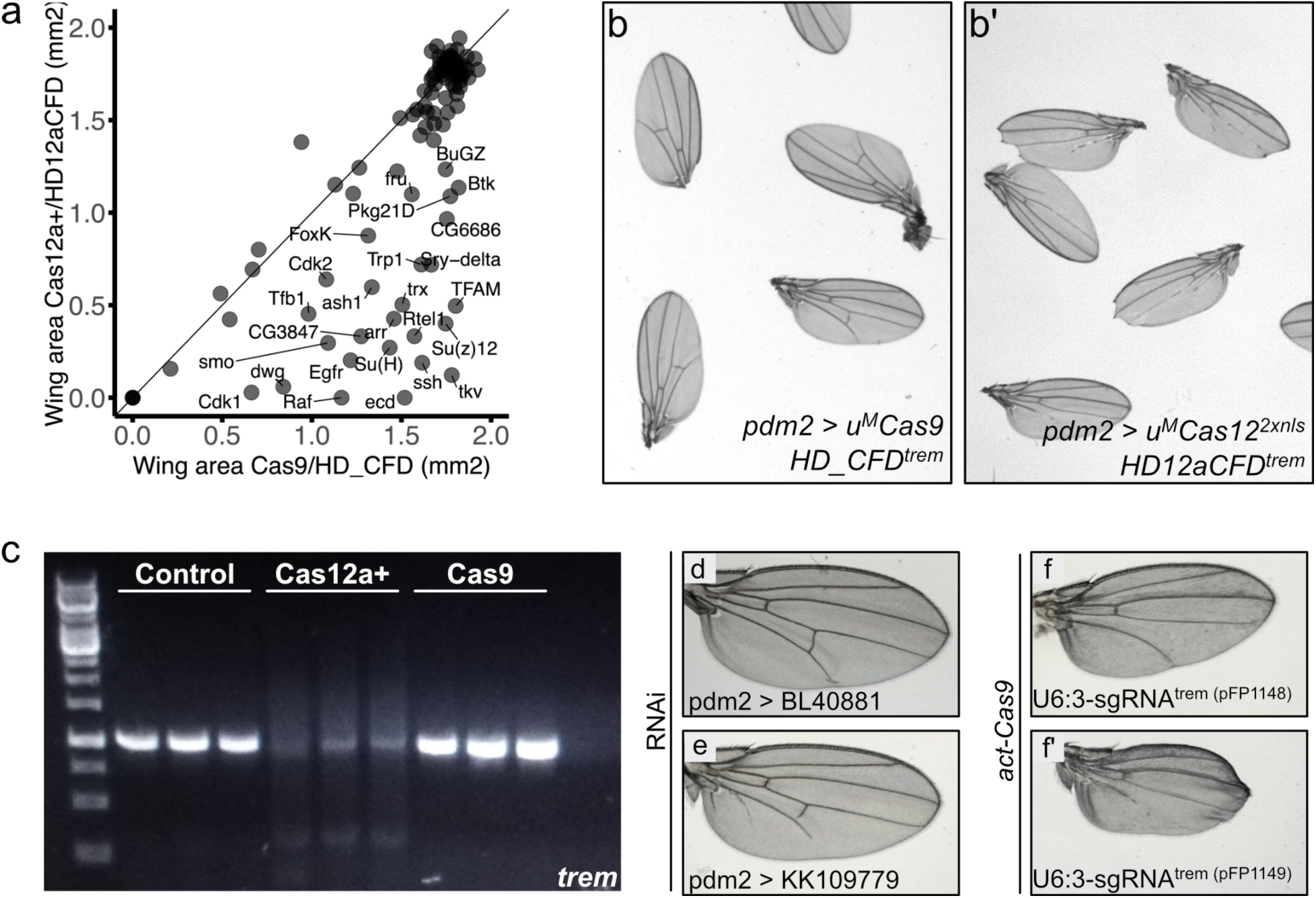
Gene disruption with HD12aCFD arrays improves recall of known and discovery of novel genes controlling wing morphogenesis. (**a**) Annotated scatter plot from Figure 3i. Several genes known to be important for wing growth show a stronger phenotype when targeted by Cas12a^+^ and HD12aCFD sgRNA arrays compared to Cas9 with HD_CFD sgRNA lines, including *tkv*, *Egfr*, *arr*, *dwg*, *ecd*, *Raf*, *Cdk1*, and *Su(h)*. (**b**) Wings from the phenotypic wing size screen presented in panel (a). Targeting *trade embargo* (*trem*) with Cas9 and a HD_CFD sgRNA pair does not lead to obvious phenotypes in the wing (b). Mutagenesis of the same gene with Cas12a+ and a HD12aCFD sgRNA array results in loss of vein tissue specifically in the posterior compartment of the wing, along with a subtle effect on shape and size of the wing (b’). Previous work suggested that *trem* does not have a role in wing morphogenesis (53). (**c**) PCR amplicons of the *trem* locus from control animals or Cas12a^+^ and HD12aCFD or Cas9 and HD_CFD sgRNA lines. Mutagenesis with HD12aCFD results in a strong reduction of amplicons of the original size, an effect that is not observed with HD_CFD. (**d**, **e**) Heterogenous effects of trem knock-down with RNAi. While RNAi line BL40881 does not result in a wing phenotype (d), knock-down with line KK109779 results in a shortening of the L5 vein and posterior cross vein (e). (**f**, **f’**) Mutagenesis with subsequently made sgRNA pairs under control of a ubiquitous *U6:3* promoter and *act-Cas9* results in lethality with occasional escapers. Escaper animals show loss of posterior veins.

